# Trailing-edge zombie forests can increase population persistence in the face of climate change

**DOI:** 10.1101/2021.12.07.471250

**Authors:** Robin R. Decker, Marissa L. Baskett, Alan Hastings

## Abstract

Climate-driven habitat shifts pose challenges for dispersal-limited, late-maturing taxa such as trees. Older trees are often the most reproductive individuals in the population, but as habitats shift, these individuals can be left behind in the trailing range edge, generating “zombie forests” that may persist long after the suitable habitat has shifted. Are these zombie forests vestiges of ecosystems past or do they play an ecological role? To understand how zombie forests affect population persistence, we developed a spatially explicit, stage-structured model of tree populations occupying a shifting habitat. Our model shows that seed dispersal from zombie forests to the range core can considerably increase the maximum rate of climate change that a population can withstand. Moreover, the entire core population can ultimately descend from recruitment-limited zombie forests, highlighting their demographic value. Our results suggest that preserving trailing-edge zombie forests can greatly increase population persistence in the face of climate change.

## Introduction

Climatic warming is expected to alter species ranges, shifting suitable habitat upslope or poleward (Walther *et al*., 2002). Many species have already shifted their ranges in response to climate change (Parmesan & Yohe, 2003), and accelerating climatic warming will force species to keep pace with shifting habitats via dispersal, adapt to new climates, or fail to persist (Loarie *et al*., 2009; Lenoir & Svenning, 2015).

Keeping pace with rapid climate change is more difficult for dispersal-limited taxa, such as plants (Corlett & Westcott, 2013; Svenning & Sandel, 2013; García *et al*., 2017; Davis & Shaw, 2001). Plant responses to climate-driven shifts in suitable habitat will depend on plant life history strategies (Aitken *et al*., 2008; Pearson *et al*., 2014; Harsch *et al*., 2016). Short-lived annual plant populations reach reproductive maturity quickly and are comprised of individuals that are approximately all the same age and undergoing the same population dynamics (Crawley, 1990). However, long-lived species, such as trees, can take decades to reach reproductive maturity and can persist for hundreds of years (Salguero-Gómez *et al*., 2016). As the climate shifts, this can create different dynamics at the leading and trailing edges of the range: many of the individuals at the leading edge may be nonreproductive, immature trees that recently colonized newly suitable habitat, while individuals at the trailing edge may be some of the oldest, largest, and most reproductive individuals, left behind in an environment that may no longer be climatically optimal (Svenning & Sandel, 2013). These mature trees would then form “zombie forests”, which may persist for decades after the climate has shifted, still dispersing seeds, many of which will likely have limited recruitment success in the suboptimal environmental conditions of the trailing edge (Kroiss & HilleRisLambers, 2015; Walck *et al*., 2011; Kueppers *et al*., 2017).

Existing theoretical frameworks provide insight into short-lived plant population dynamics in response to climate change (Potapov & Lewis, 2004; Berestycki *et al*., 2009; Zhou & Kot, 2011; Kot & Phillips, 2015; Phillips & Kot, 2015; Bouhours & Lewis, 2016), but insight into long-lived plant dynamics is sparse (but see Harsch *et al*. (2014, 2016)). Furthermore, our existing theoretical understanding of long-lived plant dynamics is based on the assumption that the environment outside of the suitable habitat is lethal (Harsch *et al*., 2016). While it is possible that no individuals survive outside of the suitable core habitat, it is more likely that at least some survive and possibly even reproduce, depending on how climatically sub-optimal environments affect plant demographic processes. For example, decreased precipitation and increased temperatures could reduce soil moisture availability (Pastor & Post, 1988). Adult plants may survive under these conditions, but reduce reproductivity, instead dedicating limited available energy towards survival and maintenance. Alternatively, survival of early developmental stages may decrease in response to climate change (Nambiar & Sands, 1993). Recent empirical studies show that forests are more likely to experience recruitment declines rather than death of adult individuals as climate warms, because early developmental stages of plants are more sensitive to climate change than mature stages (Kroiss & HilleRisLambers, 2015; Andrus *et al*., 2018; Walck *et al*., 2011; Kueppers *et al*., 2017; Davis *et al*., 2019). The way individual tree populations physiologically and demographically respond to climatically unsuitable environments could affect expectations for their ability to persist in the face of climate change.

Are zombie forests simply the living dead and remnants of ecosystems past, or do they play an ecological role in supporting population persistence in the face of climate change? We still do not fully understand the demographic role that each edge of the range plays in promoting the establishment of forests as they shift to higher latitudes and upslope. The leading range edge has been identified as a conservation priority, because it shifts into climatically favorable environmental conditions and therefore could have a naturally higher survival rate (Gibson *et al*., 2009; Rehm *et al*., 2015). However, zombie forests in the trailing range edge can produce and disperse many seeds, which could increase population growth throughout the entire range. Zombie forests thereby could potentially play a key role in increasing population persistence in the face of climate change, despite their location in the suboptimal environmental conditions of the trailing range edge. Identifying whether any demographic characteristics of zombie forests promote population persistence could help distinguish the cases in which preservation of the trailing range edge benefits population range shifts and persistence.

To elucidate whether and under what conditions zombie forests play a role in climate-driven tree range shift dynamics, we develop a stage-structured spatial integrodifference model that accounts for individuals left behind as suitable habitat shifts. With this model, we explore the following questions: (1) Do zombie forests influence population persistence under climate change and, if so, how? (2) How does dispersal ability mediate the response of the population to climate-driven range shifts? (3) How do the answers to these questions depend on the effect of climate change on tree survival, growth, and reproduction? (4) What mechanisms explain the effect of zombie forests on the rest of the population?

## Methods

### Model description

The purpose of the model is to determine how mature trees left trailing behind a moving habitat patch affect the population’s ability to persist in response to climate change. We model a habitat patch that shifts through continuous space over discrete time to simulate a climate-driven niche shift. Population growth dynamics are stage-structured and independently vary behind the patch, within the patch, and in front of the patch. Seed dispersal is symmetric and average dispersal distances are invariable throughout the tree’s range (fig. 1).

**Figure 1:**
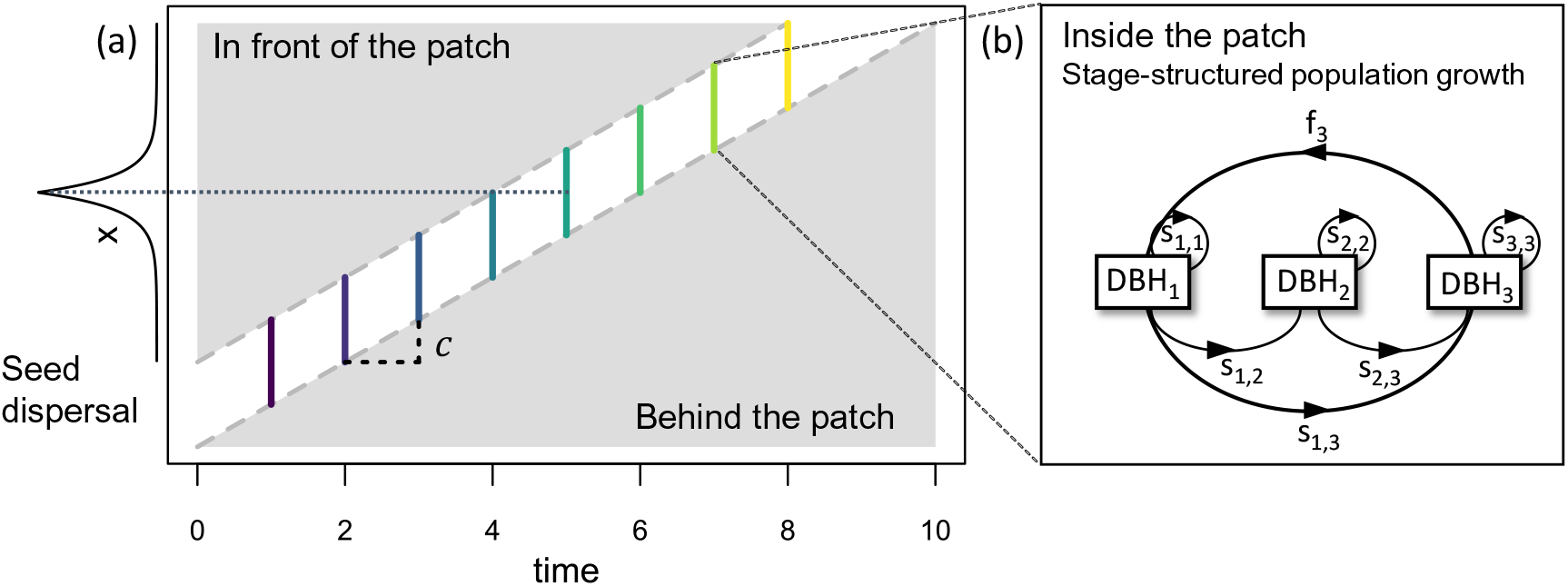
Moving habitat patch model overview. (a) Climate change at rate *c* causes the habitat patch to shift in space *x* each season. In the null model, growth is impossible in the environments in front of and behind the patch. We independently vary the environments in front of and behind the patch, testing four different environmental scenarios (see table 1). (b) Inside the shifting habitat patch, the stage-structured tree population grows and disperses seeds following a Laplace dispersal kernel. Population growth follows a stage-structured model with size classes based on trunk diameter at breast height (DBH); three size classes in the case of sugar maple. Transitions *s_i,j_* represent survivorship or growth from stage *i* to stage *j*, and transitions *f_i_* represent reproductive transitions from stage *i* to the seedling stage.

Because trees often reproduce annually and have well-defined growth and seed dispersal stages, we use integrodifference equations to model these populations. Integrodifference equations are discrete-time, spatially explicit models that track the continuous distribution of population density over space. Integrodifference equations are commonly used to model the spatial spread of invading organisms (Kot *et al*., 1996; Cuddington & Hastings, 2004; Hastings *et al*., 2005) and plant population range shifts in response to climate change (Zhou & Kot, 2011; Harsch *et al*., 2014; Kot & Phillips, 2015; Phillips & Kot, 2015; Bouhours & Lewis, 2016; Harsch *et al*., 2016).

We modeled the population density of each tree developmental stage or size class at any spatial location *x* and any discrete time *t* as the vector **N***_t_*(*x*), where each entry in the vector corresponds to a developmental stage. Each time step *t*, population growth and dispersal occur. The transition matrix **A** describes the probabilities of transition between stages due to survivorship and growth, as well as stage-specific fecundities (Caswell, 2001). The matrix-valued function **H**(**A***, x, t*) takes as an input the matrix **A** and produces a matrix that is obtained by modifying **A** spatially to produce a new matrix to delineate the patch boundaries and control patch movement with climate change. A matrix of dispersal kernels **K**(*x − y*) describes the seed dispersal stage. Therefore, the age-structured growth-dispersal shifting habitat model has the general form:

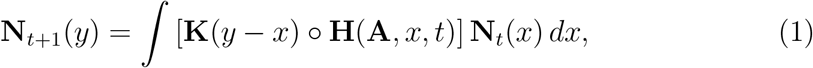

where *◦* is the Hadamard product (Caswell, 2019), element-by-element multiplication of the matrices **K** and **H**(**A***, x, t*).

Each element *k_ij_* (*y − x*) of the dispersal kernel matrix **K**(*y − x*) is a function describing how individuals in stage *j* disperse to stage *i*, from any location *x* to any location *y*. For transitions associated with survivorship or growth (e.g. growth from a medium diameter tree stage to a large diameter stage), there is no movement of the individual tree, and the corresponding entry in the dispersal kernel matrix describes the stationary transition (using the Dirac delta function (Harsch *et al*., 2014)). For transitions associated with seed production, we model seed dispersal with a Laplace distribution kernel, with parameter *b*, which represents the mean dispersal distance of seeds (Kot *et al*., 1996):

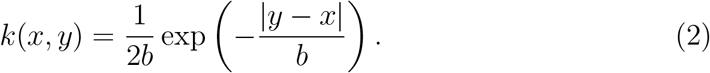

To model a changing climate, we use the function **H**(**A***, x, t*) to delineate a patch of length *L* centered at the origin, which increases by *c* units each year, where *c* is the speed of climate change. The transition matrix **A** governs the population dynamics inside the patch, and a matrix of modifiers **M**, with the same dimensions as **A**, modifies the population dynamics outside of the patch to model the reduction in fecundity, recruitment, survivorship, or growth associated with the suboptimal environment outside of the patch. If all entries in **M** are 1, then the environment outside of the patch will be identical to the environment inside of the patch. If all entries in **M** are 0, then the environment outside of the patch will be lethal. The matrix of modifiers **M** varies independently behind and in front of the habitat patch. If **M***_b_* is the matrix of modifiers for behind the patch, and **M***_f_* is the matrix of modifiers for in front of the patch, then

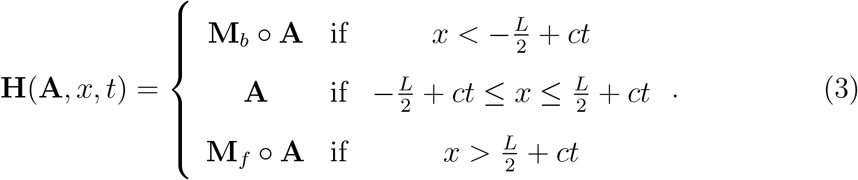

The general form of **A** with *n* size classes, transitions *s_i,j_* that represent survivorship or growth from stage *i* to stage *j*, and transitions *f_i_* that represent reproductive transitions from stage *i* to the seedling stage is then

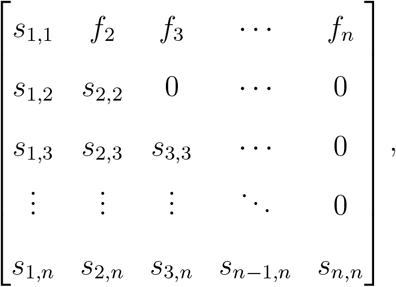

and the general from of the matrix of modifiers **M** with recruitment modifier *m_r_*, fecundity modifier *m_f_*, and survivorship and growth modifier *m_s_* is

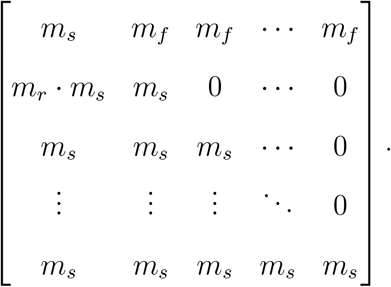

### Model parameterization and tree species characteristics

We consider two species of North American trees which may face climate-driven habitat shifts in the near future: sugar maple (*Acer saccharum*), a deciduous species whose range broadly covers most of the northeast United States, and white fir (*Abies concolor*), a coniferous species which is distributed across the western United States, with large populations in California. Of east coast tree species, sugar maple is one of the most vulnerable to climate change (Rogers *et al*., 2017), and its area of suitable habitat could shift north more than 20 km over the next century (Iverson & Prasad, 2002). White fir has been long considered much less vulnerable, but severe drought in California, which is predicted to increase in frequency (Ummenhofer & Meehl, 2017), has increased white fir mortality (Guarín & Taylor, 2005), and climate change is projected to decrease white fir productivity and further increase susceptibility to mortality (Battles *et al*., 2007).

We retrieved transition matrices for sugar maple and white fir from COMPADRE, the plant matrix database (Salguero-Gómez *et al*. (2015), see Appendix S1 in Supporting Information). The sugar maple transition matrix models the population dynamics of an old growth sugar maple forest located in Brownfield Woods, Illinois in an area that experienced little disturbance and low neighbor density (Lin & Augspurger, 2008). Lin & Augspurger (2008) used historical tree maps based on a census period from 1951-1988 to calculate the transition probabilities for the three size classes. The white fir transition matrix models the population dynamics of a white fir-dominated area located in Hodgdon Meadow in Yosemite National Park, California (Van Mantgem & Stephenson, 2005). Van Mantgem & Stephenson (2005) calculated the transition probabilities based on a census period from 1991 to 1994 in an unlogged area of forest.

For parameterizing the dispersal kernel, sugar maple is a wind-dispersed species, with nearly all seed dispersal occurring within 15 m of the forest edge (Hughes & Fahey, 1988). Although some seeds will disperse 15 m or greater from the parent tree, average seed dispersal distance is likely much lower. We tested a range of average dispersal distances from 0 m to 20 m for sugar maple. White fir seeds are wind-dispersed from the cone as it falls, with nearly all seeds landing in an opening 1.5 to 2 times the height of the tree (Zouhour, 2001). The radius of seed dispersal is half this opening (.75 to 1 times the height of the tree), but most seeds land much closer to the tree, especially when wind speeds are low. While white fir in California can reach heights of 40 to 55 m (Jones, 1974), the average height of any given mature tree is considerably less. We therefore tested a range of average dispersal distances from 0 m to 7 m for white fir. We chose patch widths of 10 m and 1 m for sugar maple and white fir, respectively, which are similar to values used commonly in the moving habitat model literature (Harsch *et al*., 2016), and are scaled appropriately to the range of mean dispersal distances tested. See Appendix S1 for transition and dispersal matrices.

### Model analyses

We used the critical speed of climate change as a measure of a population’s ability to persist under climate change. The critical speed of climate change is the maximum speed of directional climate change, measured in meters per year, that allows a population to persist long-term without extirpation (fig. 2). Speeds of climate change above the critical speed cause the population to eventually decline to zero (Zhou & Kot, 2011). We numerically calculated the critical speed of climate change by running simulations under a range of values for *c*, and determined population persistence by whether the population increased or declined after 100 years of climate change.

**Figure 2:**
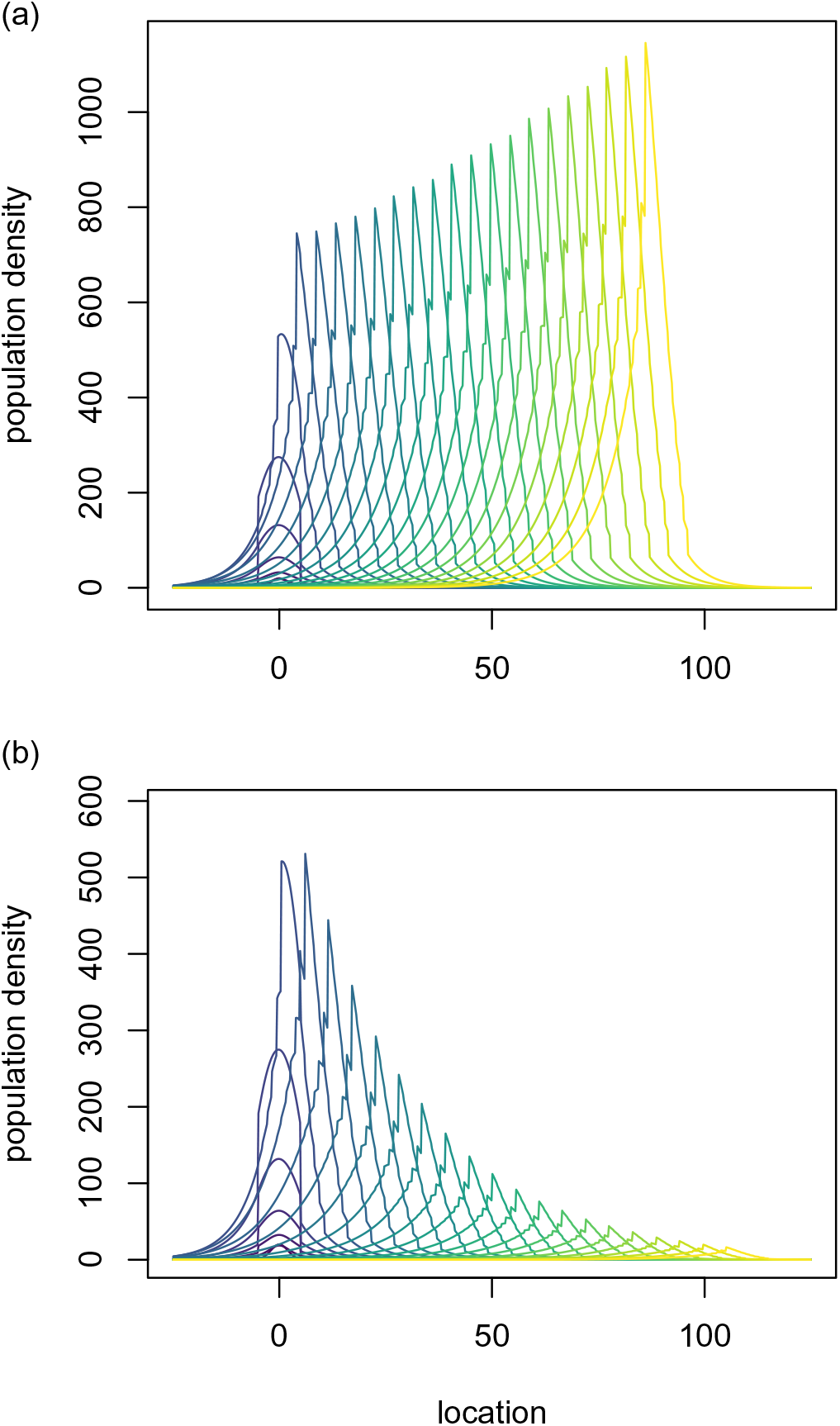
Illustration of the critical speed of climate change and typical model output. In both panels, a sugar maple population with a patch size of 10 m, average dispersal distance of 5 m, reduced survivorship and growth (environmental scenario 4) behind the patch, and no survival (environmental scenario 1) in front of the patch grows for 25 generations without climate change, followed by 100 generations with climate change. Each curve represents the population density distribution every 5 generations, with early generations in blue and later generations in yellow. The critical speed of climate change for this population is 0.92 m/year. The population grows when *c* = 0.91 (a) and fails to persist when *c* = 1.1 (b).

To determine how zombie trees behind the patch influence the critical rate of climate change, we numerically computed and compared the critical speed of climate change in populations with zombie forests (scenarios 2-4, table 1) to populations without zombie forests (scenario 1, table 1). Because the effect of climatic warming on tree demographic processes is not yet completely understood, we compared three different environmental scenarios that represent possible ways that climate change could affect tree demographics (table 1). These included recruitment failure (scenario 2), no seed production (scenario 3), and reduced survivorship and growth (scenario 4). The long-term population growth rate (the dominant eigenvalue, *λ*) in the unmodified transition matrix was 1.27 for the sugar maple population, and 1.09 for the white fir population. In the fatal environment (scenario 1), the dominant eigenvalue was 0. In the other scenarios, the dominant eigenvalue was 0.81 for sugar maple, and 0.97 for white fir.

**Table 1:** Environmental scenarios tested outside the patch and corresponding values for the recruitment *m_r_*, fecundity *m_f_* and survivorship/growth *m_s_* modifiers. Note that the unmodified transition matrix is used only inside the moving habitat patch.

| Environmental Scenario | Description | $m_r$ | $m_f$ | $m_s$ |
| --- | --- | --- | --- | --- |
|  | Unmodified transition matrix **A** | 1 | 1 | 1 |
| 1 | Fatal environment | 0 | 0 | 0 |
| 2 | No recruitment | 0 | 1 | 1 |
| 3 | No fecundity | 1 | 0 | 1 |
| 4 | Reduced survivorship & growth | 1 | 1 | <1 |

We explicitly explored the role of zombie forests by testing these environmental scenarios exclusively in the area behind the moving habitat patch. For completeness, in Appendix S4, we also separately tested the same scenarios in the area in front of moving habitat patch and simultaneously both behind and in front of the moving habitat patch. To determine the influence of dispersal ability, we repeated these calculations across a range of ecologically reasonable values for mean seed dispersal distances (Hughes & Fahey, 1988; Zouhour, 2001; Jones, 1974; Harsch *et al*., 2016).

To determine whether zombie forests contribute to population persistence, we compared the critical rate of climate change in scenario 1 to the other scenarios. To understand the mechanism by which zombie forests aid the core population, we computed the proportion of the core population that descended from zombie forests over time in each of the four environmental scenarios (Appendix S2). We also computed the proportion of the population comprised of zombie individuals over time for each environmental scenario. In these cases, we assumed dispersal ability was intermediate, with an average dispersal distance of 5.0 m for sugar maple and 3.0 m for white fir. If small proportions of the population descend from zombie forests, but a large proportion of the whole population is the zombie forest, then zombie forests help the population simply by surviving and growing. Alternatively, if large proportions of the population descend from zombie forests over time, then zombie forests help the population by dispersing seeds into the population core.

We solved the model in R version 4.0.3 (R Core Team, 2021) using R’s built-in integration routine, which uses globally adaptive interval subdivision in connection with extrapolation by Wynn’s Epsilon algorithm, with Gauss–Kronrod quadrature as the basic step (Piessens *et al*., 1983). We initialized the model with a small, normally distributed population of seedlings, centered in the middle of the patch with standard deviation 2.0. For each model run, the population grows for 50 generations without climate change to ensure consistent population growth over space (Appendix S3).

## Results

Explicitly including individuals at the trailing edge of the patch increased the critical speed of climate change that a population could endure (figure 3). Across all environment scenarios, populations with shorter-distance seed dispersal had correspondingly lower critical speeds of climate change. Intermediate dispersal ability, relative to patch size, increased the rate of climate change a population could withstand. However, high average dispersal distances relative to patch size decreased a population’s ability to persist in high speeds of climate change. Populations with zombie forests in a state of recruitment failure (environmental scenario 2) had the highest critical rates of climate change. Populations with zombie forests with reduced survivorship and growth (scenario 4) had higher critical rates of climate change than populations without zombie forests (scenario 1). Populations with zombie forests that produced no seeds (scenario 3) had nearly the same critical rates of climate change as populations with no zombie forests (scenario 1). In contrast to trailing-edge populations, adding leading-edge populations increased the critical rate of climate change for scenario 3 more than scenarios 1, 2, and 4 (Appendix S4).

**Figure 3:**
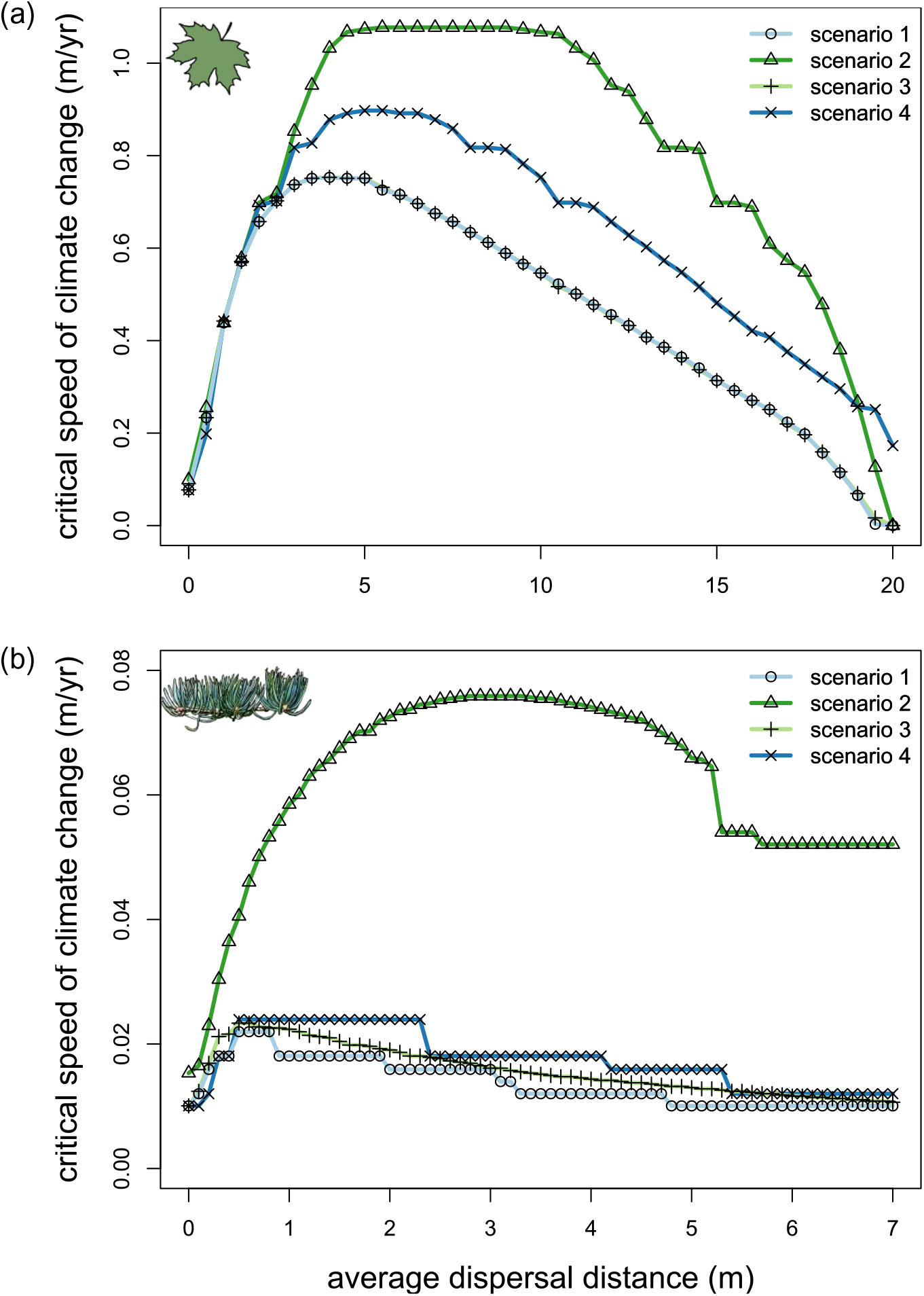
Effect of environment behind the patch and dispersal distance on the critical speed of climate change for population persistence in sugar maple (a) and white fir (b), with no survival (environmental scenario 1) in front of the patch. Behind the patch environmental scenarios tested include no survival (scenario 1), no recruitment (scenario 2), no fecundity (scenario 3), and reduced survivorship and growth (scenario 4).

When zombie forests were present and producing seeds (scenarios 2 and 4), the proportion of the core population that descended from zombie forests increased over time (fig. 5). For both species tested, virtually all of the core population descended from the zombie forest by 350 years. By contrast, when zombie forests faced death in the rear range edge or did not produce seeds (scenarios 1 and 3), none of the population descended from the zombie forest. The proportion of the population comprised of zombie trees varied between species tested and among different environmental scenarios. The fir population had roughly twice as many zombie trees as the maple population across all environmental scenarios. When zombie forests did not produce seeds (scenario 3), they comprised a bigger proportion of the population than when they did produce seeds (scenarios 2 and 4). Even when zombie trees died at the end of the year after entering suboptimal habitat at the rear range margin (scenario 1), zombie trees still comprised a small proportion of the population each year.

**Figure 4:**
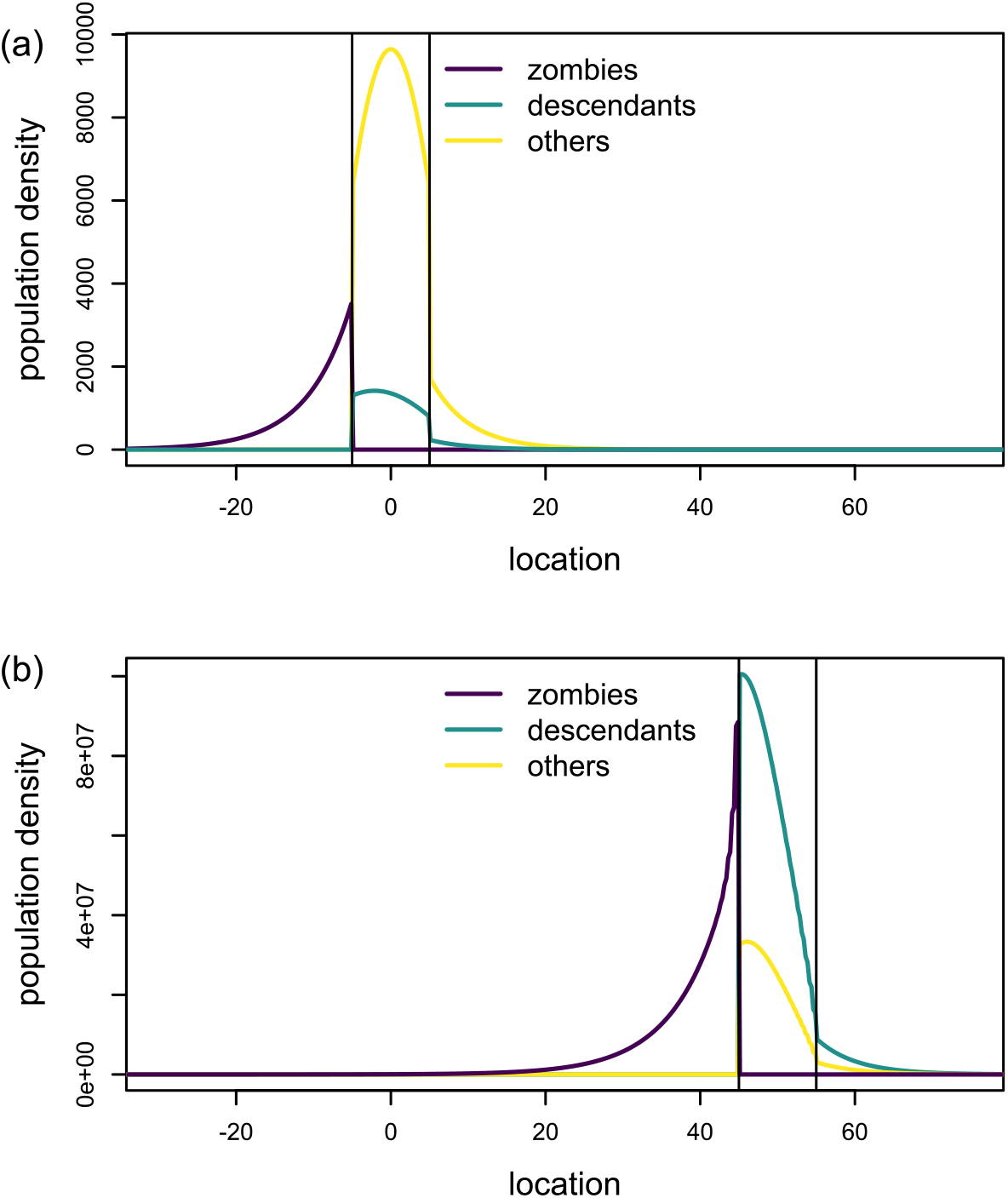
Illustration of zombie forests and their descendants, before (a) and after (b) climate change. Black vertical lines indicate the boundaries of the moving habitat patch. Distributions of the tree population behind the patch (i.e. zombies), the tree population descended from zombies, and all other individuals in the population are show in purple, teal, and yellow, respectively. In front of the patch, there is no survival (environmental scenario 1), but behind the patch, zombie forests grow with reduced survivorship and growth (scenario 4). (a) A sugar maple population grows for 50 generations without climate change. (b) After 100 generations of climate change, a larger proportion of the population is descended from zombies.

**Figure 5:**
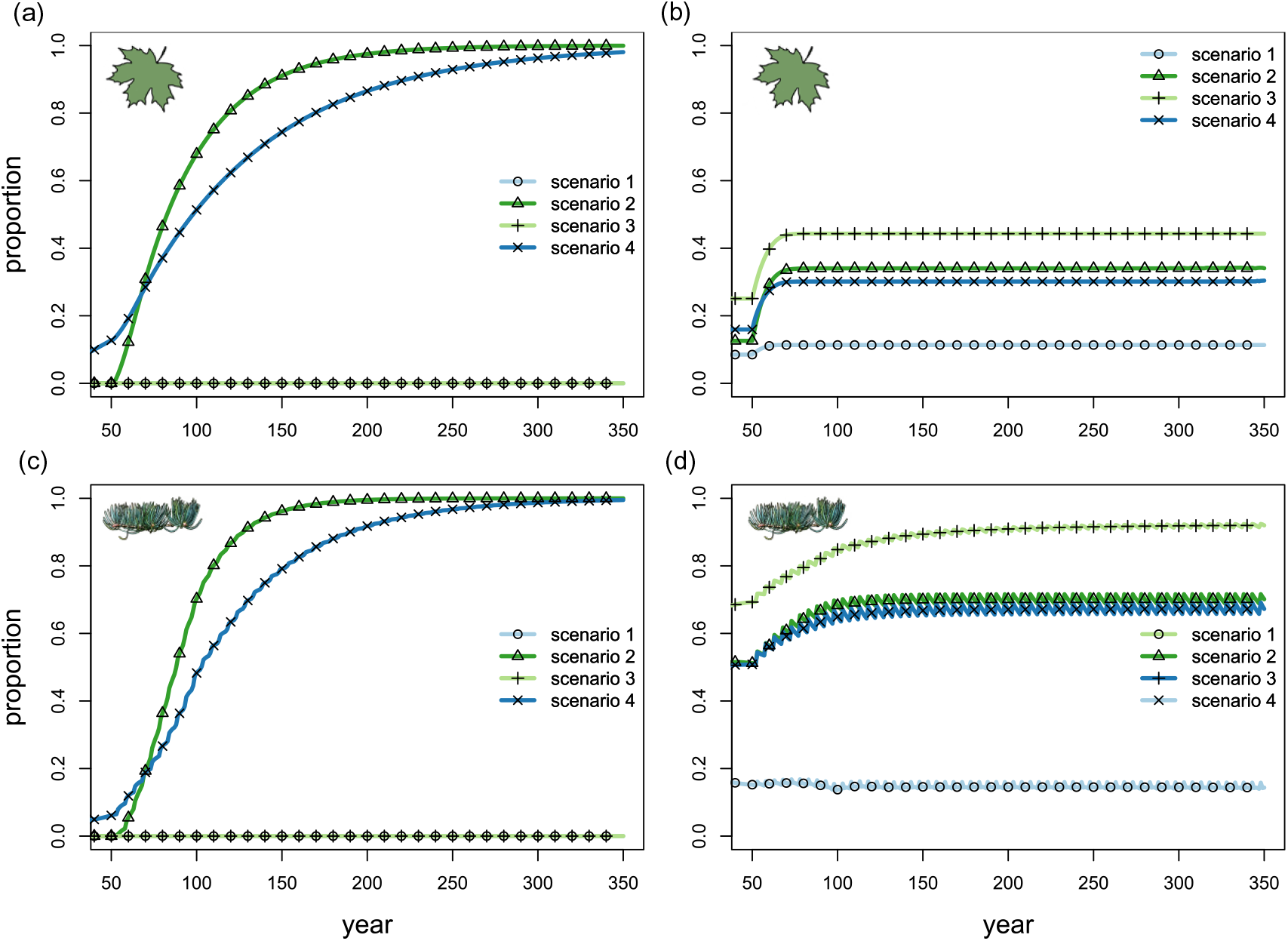
Proportion of the core population descended from the zombie forest over time for (a) sugar maple and (c) white fir populations with no survival (environmental scenario 1) in front of the patch. Proportion of the forest that is comprised of zombie individuals for (b) maple and (d) fir. Behind the patch environmental scenarios tested include no survival (scenario 1), no recruitment (scenario 2), no fecundity (scenario 3), and reduced survivorship and growth (scenario 4). Both populations had intermediate average dispersal distances, with maple 5 m and fir 3 m, and tolerable rates of climate change, with maple 0.5 m/yr and fir 0.02 m/yr, which started at year 50.

## Discussion

The presence of trailing-edge zombie forests can substantially increase the maximum rate of climate change a population can tolerate (fig. 3). Our model suggests that zombie forests must have both high amounts of seed production and high survival to protect the population from rapid climate change, because zombie forests supply propagules to the core population. In contrast, high rates of growth or recruitment in zombie forests do not improve the population’s response to elevated rates of climate change. Note that the opposite is true for leading-edge forests: they contribute most when the environment promotes high rates of growth and survival for seedling stages (Appendix S4). Therefore, the demographic characteristics of zombie forests that provide the greatest increases in the critical speed of climate change match the empirically determined characteristics of forests stressed by climate change (Walck *et al*., 2011; Kueppers *et al*., 2017). This suggests that zombie forests might have considerable potential to protect entire tree populations from rapid rates of climate change.

When mature trees within the zombie forests survive and reproduce, the entire core population ultimately descends from the zombie forest (fig. 5a,c), suggesting that zombie forests can become increasingly important over time. This somewhat counter-intuitive pattern is explained by the typical life cycle of an individual tree in a shifting habitat. As the suitable habitat shifts, the core population that occupied the previously suitable habitat is left behind, and becomes the zombie forest. Zombie trees at the back of the patch then initiate a cascade of events: they produce a large number of seeds, a proportion of which then land in the suitable habitat and form the core of the population, which matures over time and becomes the next generation of zombie forest as the suitable habitat shifts. The cycle repeats driven by the dynamics of zombie trees at the trailing edge, which sustain the shifting population. If the trailing edge and core populations were separated from each other, neither would persist in the presence of moderate to high rates of climate change (fig. 3). However, they persist together, in part because the zombie forest acts as a seed source while the core acts a seed sink (Watkinson & Sutherland, 1995). The value of the trailing edge as a source of genetic diversity is well-recognized (Hampe & Petit, 2005; Provan & Maggs, 2012; Rehm *et al*., 2015), but our results suggest that the trailing edge can also play an important demographic role in supporting population persistence.

Zombie forests can also play a role in maintaining the population in front of the patch. The core population, which is partly produced by the zombie forest, matures and seeds the area in front of the patch. If recruitment in front of the patch is high enough (fig. S3), these seedlings in front of the patch mature and eventually become the core population as the area of suitable habitat shifts in response to climate change. The model suggests that there may be minimal benefit to increasing fecundity (fig. S3) in front of the patch. High survival and growth at the front of the patch, especially for the youngest seedling size class, are necessary to increase the critical speed of climate change.

### The role of dispersal

The benefit of zombie forests hinges on the population having an intermediate average dispersal distance (figure 3). Zombie forests may not help tree populations that are dispersal limited. At low average dispersal distances, seeds may not disperse far enough in front of the suitable habitat to keep up with high climate-driven habitat shift rates. However, extremely high average dispersal distances may not be beneficial either, causing seeds to disperse very far behind or in front of the suitable habitat (Zhou & Kot, 2011), which decreases the proportion of the population occupying the optimal environment inside of the suitable habitat. Populations with larger habitats (figure 3a) can persist under higher rates of climate change and have maximum critical speeds of climate change at higher dispersal levels than populations on smaller habitat patches (figure 3b). This result is rooted in the theory of dispersal success in a stationary habitat patch (Van Kirk & Lewis, 1997) and agrees with the findings of previous moving habitat model studies (Potapov & Lewis, 2004; Berestycki *et al*., 2009; Zhou & Kot, 2011; Harsch *et al*., 2016). Our results extend this theory to explicitly consider the role of the environment outside the area of suitable habitat.

### Species-specific impacts of zombie forests

We focused our analysis on two tree populations: a deciduous species with a high population growth rate and a coniferous species with a low population growth rate. Zombie forests had a similar qualitative effect on both of these species in our analysis (figs. 3, 5, S3, S4), but there were several key differences that can be attributed to the differences in population growth rate. Our model suggests that populations with lower growth rates can have correspondingly lower critical speeds of climate change, extending the findings from other moving habitat models (Zhou & Kot, 2011; Harsch *et al*., 2014, 2016) to account for the role of zombie forests. However, zombie forests can have a proportionally greater influence on slow-growing populations: recruitment-limited zombie forests with intermediate dispersal ability increased the maximum tolerable speed of climate change by 377% in the slow-growing population as compared to 43% in the fast-growing population. Survival of older trees in the back part of the range can help slow-growing populations by providing them more time to reach reproductive maturity: individual trees spend the early stages of their lives in the core of the range, and the later stages of their life in the trailing range edge as the range shifts. The importance of this mechanism is evidenced by the observation that most individuals are members of the zombie forest in the slow-growing population, which is not the case for fast-growing population (fig. 5b,d).

### Assumptions, simplifications and future directions

While tree demographic information is readily available for a variety of species, it is much more difficult to find empirical characterizations of seed dispersal kernels. We therefore assumed that both the sugar maple and white fir dispersal kernels generally followed a Laplace distribution, and that mean dispersal distances fell between 0 and 15 m for sugar maple and between 0 and 7 m for white fir, based on generalizations from the literature. Across this range of values, the critical speed of climate change varied considerably. Empirically describing, generalizing and predicting dispersal processes in tree populations is critically necessary in order to understand how tree populations will respond to climate change (Aslan *et al*., 2019; Beckman *et al*., 2020). For instance, it may be the case that larger mature trees generally disperse seeds farther than smaller mature trees, particularly for species like white fir where dispersal radius depends on tree height (Zouhour, 2001). This could increase the importance of zombie trees at the trailing range edge, as more of their seeds could land within the ideal habitat patch. We also assumed constant dispersal in time, but the importance of zombie forests could change with temporally variable dispersal (Ellner & Schreiber, 2012; Williams & Hastings, 2013). Such variation in seed dispersal might arise from stochastic variation in wind, for example in the case of wind-dispersed species such as sugar maple and white fir (Nathan & Muller-landau, 2000). Because variable dispersal can increase population spread rates and persistence in metapopulations (Ellner & Schreiber, 2012; Williams & Hastings, 2013), variability in dispersal could increase the critical rate of climate change in range-shifting populations. This could decrease the reliance of populations on zombie forests at intermediate rates of climate change, but increase the importance of zombie forests at high rates of climate change.

We additionally made several simplifying assumptions about the structure of the model. We ignored the possibility of evolution, which could allow species to adapt to climate change, thereby minimizing the rate of shift necessary to keep pace with climate change (Nadeau & Urban, 2019; Diamond, 2018; Thurman *et al*., 2020). However, the spatial population genetics of shifting populations could cause movement of seeds from the trailing edge and from the patch center to the leading edge. This could genetically swamp the populations at the leading edge, limiting the possibility of adaptation to changing climatic conditions and increasing the population’s dependence on dispersal ability to persist in the face of climate change (Bridle & Vines, 2007; Duputié *et al*., 2012). Evolution of dispersal ability adds complexity to these dynamics, and could ultimately accelerate expansion of populations into newly suitable habitat (Block & Levine, 2021), especially with the aid of zombie forests increasing population seed production.

We assumed that the population shifts into an environment with no anthropogenic barriers (Travis, 2003; Littlefield *et al*., 2019; Musgrave & Lutscher, 2014), no topographical complications (Elsen *et al*., 2020; Ackerly *et al*., 2020), and no biotic interactions (Urban *et al*., 2012; Usinowicz & Levine, 2018; Brooker *et al*., 2007). Competitive interactions between shifting plant communities could hinder the ability of zombie forests to increase population persistence, especially if other plant communities shift into the area occupied by the zombie forests and compete with these trees for limited resources. However, zombie forests could also support the core area of the range by increasing the dispersal pressure of the population as it collides with preexisting communities (Wallingford *et al*., 2020). In either case, explicitly including the leading and trailing range edges in range shift models, as we have done here, is a step in the direction of understanding how colliding communities could respond to climate-driven range shifts.

### Management Implications

The idea that climate-induced range shifts could create stands of zombie forests raises the question of if there is any point in preserving these trees that are destined to die anyway as their species’ range shifts progressively away. At the same time, there is substantial debate about the optimal strategies to preserve forests struggling to keep pace with rapid climate change (Millar *et al*., 2007; Dawson *et al*., 2011; Heller & Zavaleta, 2009). One proposed strategy is to preserve individuals at the leading edge of the range (Gibson *et al*., 2009; Rehm *et al*., 2015), which our model suggests are often seedlings and young trees. However, in practice, increasing seedling and young tree survival at the leading range edge until environmental conditions improve may not be possible, because decades may pass before conditions improve and seedlings have naturally high mortality rates due to competition and herbivory (Collet & Le Moguedec, 2007; Keeton, 2008). Additionally, seedling mortality may be caused by aspects of the environment that are difficult to control, such as fluctuations in temperature combined with low soil moisture availability (Niinemets, 2010). In species where the primary impact of climate change is decreased seed production at the leading range edge, but otherwise high seedling survival, protecting the leading edge can greatly increase population persistence. However, in species where climate change limits recruitment at the trailing range edge, it might be more important to protect zombie forests, especially by increasing dispersal from the trailing edge to the core population. Therefore strategies that promote seed dispersal, such as removing anthropogenic barriers (Caplat *et al*., 2016), particularly in animal-dispersed plant populations (Suárez-Esteban *et al*., 2013) and especially near the rear edge boundary, may help populations that otherwise have naturally high average dispersal distances. Our model shows that these strategies may not be as effective for weakly-dispersing populations, and poor dispersers may therefore require more intensive management strategies, such as managed relocation (Schwartz *et al*., 2012; Gallagher *et al*., 2015; Koralewski *et al*., 2015), to keep pace with changing climate.

## Acknowledgements

This work was supported by the Institute for the Study of Ecological and Evolutionary Climate Impacts (ISEECI) funded through the University of California Office of the President Multicampus Research Programs and Initiatives, Award #CA-15-328887, a National Science Foundation Graduate Research Fellowship to R.R.D. (NSF award no. 1650042), NSF grant no. DEB-1655475 to M.L.B., and NSF grant no. DMS-1817124 to A.H..

## Appendix S1: Tree transition and dispersal matrices

The sugar maple population is divided into three size classes based on trunk diameter at breast height (DBH): individuals *≤* 7.6 cm, individuals between 7.7 and 19.9 cm, and individuals between 20 and 39.1 cm. The transition matrix for sugar maple (Salguero-Gómez *et al*., 2015; Lin & Augspurger, 2008) inside the patch is

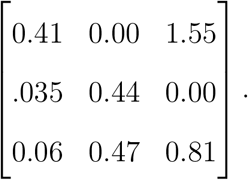

The white fir population is divided into five size classes also based on DBH: individuals between 0.0 and 0.5 cm, individuals between 5.1 and 10.0 cm, individuals between 10.1 and 20.0 cm, individuals between 20.1 and 40.0 cm, and individuals larger than 40.0 cm. The transition matrix for white fir (Salguero-Gómez *et al*., 2015; Van Mantgem & Stephenson, 2005) inside the patch is

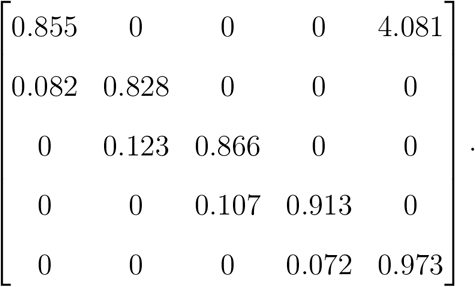

The dispersal kernel matrix for sugar maple is

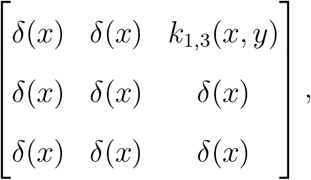

and the dispersal kernel matrix for white fir is

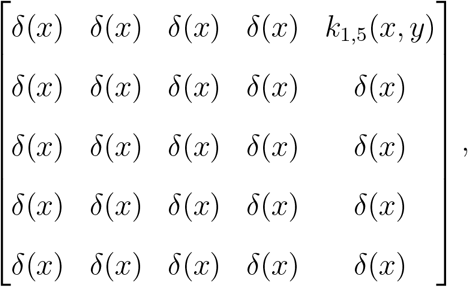

where *δ*(*x*) is the Dirac delta function, and 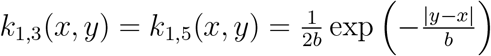, with *b* ranging from 0 to 20 m for sugar maple and from 0 to 7 m for white fir, as described in the text.

## Appendix S2: Computing the proportion of the core population descended from zombie forests

We partitioned the total population, **N***_t_*, into three components: (1) the zombie forests, which are individuals trailing behind the suitable habitat, **N***_z,t_*, (2) the descendants of the zombie forests in the core or leading range edge, **N***_d,t_*, and (3) all other individuals, which are trees that are in the core or leading range edge, but do not have a zombie ancestor, **N***_o,t_*, such that

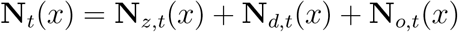

for all values of *x*.

Then, at any time *t*, the proportion of the core population descending from the zombie forest is

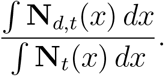

Similarly, the proportion of the population comprised of zombie individuals is

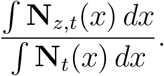

## Appendix S3: Equilibrium population density

Population growth is not density dependent in the model. Population density therefore never reaches a constant distribution over space, but the population eventually grows everywhere at all points in space. This is preceded by a period of transient population dynamics, where the population grows at some points in space, but declines at other points in space, based on the initial population density distribution. Table S1 lists the first generation that the population grows at every point in space, for each environmental scenario, each habitat configuration, and each species. In each case, we allow the model to run for 50 generations without climate change before initiating climate change. This initial population growth period is long enough to ensure consistent growth dynamics throughout the habitat. In all cases, populations reach a pattern of sustained growth by 20 generations, for both sugar maple (fig. S1) and white fir (fig. S2).

**Table S1:** The first generation at which the population grows at every point in space. Habitat configuration describes which areas of the habitat correspond to nonzero transition matrices and are thereby capable of supporting population growth: either the area behind the traveling habitat patch or the area in front of the traveling habitat patch, or both. Environmental scenario describes which transition matrix is used in the back and/or front of the traveling habitat patch: (1) the zero matrix (death outside the patch), (2) no recruitment, (3) no fecundity, or (4) reduced survivorship and growth.

| Species | Configuration | Scenario | Generation |
| --- | --- | --- | --- |
| sugar maple | back and front | 1 | 20 |
| sugar maple | back and front | 2 | 20 |
| sugar maple | back and front | 3 | 20 |
| sugar maple | back and front | 4 | 16 |
| sugar maple | back | 2 | 20 |
| sugar maple | back | 3 | 20 |
| sugar maple | back | 4 | 18 |
| sugar maple | front | 2 | 20 |
| sugar maple | front | 3 | 20 |
| sugar maple | front | 4 | 18 |
| white fir | back and front | 1 | 20 |
| white fir | back and front | 2 | 20 |
| white fir | back and front | 3 | 20 |
| white fir | back and front | 4 | 18 |
| white fir | back | 2 | 20 |
| white fir | back | 3 | 20 |
| white fir | back | 4 | 19 |
| white fir | front | 2 | 20 |
| white fir | front | 3 | 20 |
| white fir | front | 4 | 19 |

**Figure S1:**
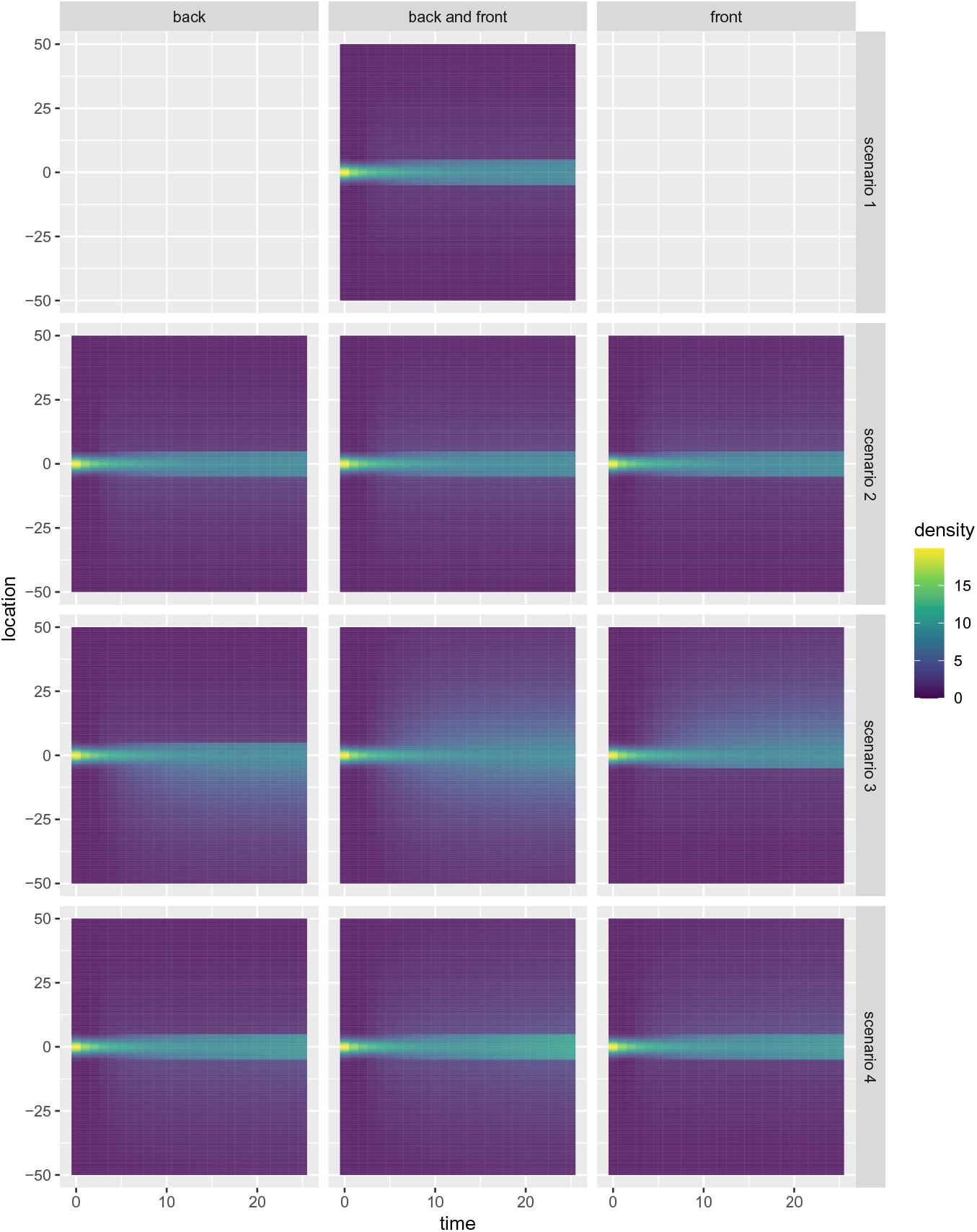
Sugar maple population density over space and time for each habitat configuration and environmental scenario. Habitat configurations include back, front, or both back and front of the traveling habitat patch, indicating which areas of the habitat are capable of supporting population growth. Environmental scenario describes which transition matrix is used in the back and/or front of the traveling habitat patch: (1) the zero matrix (death outside the patch), (2) no recruitment, (3) no fecundity, or (4) reduced survivorship and growth. In all cases, populations reach a pattern of sustained growth by 20 generations.

**Figure S2:**
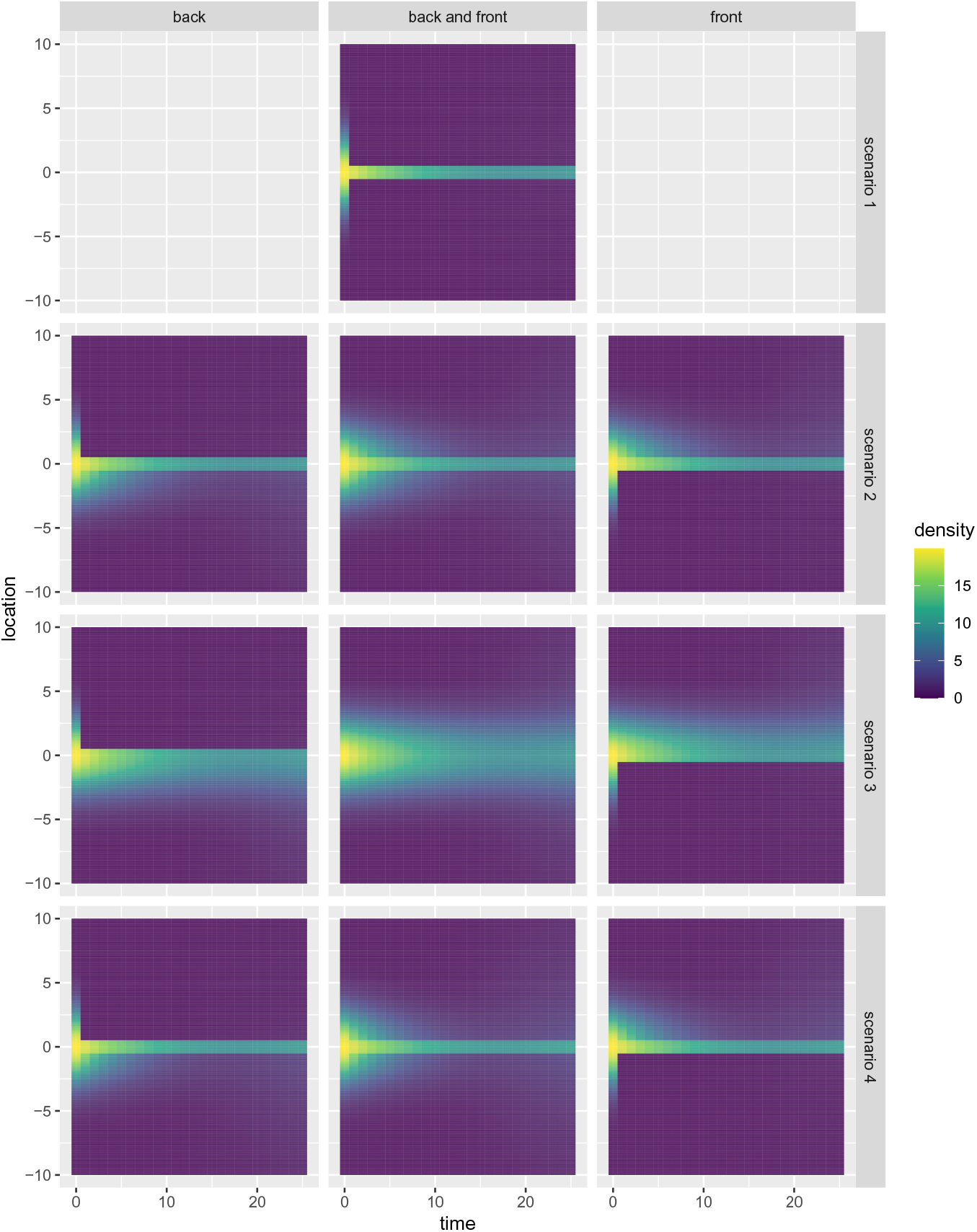
White fir population density over space and time for each habitat configuration and environmental scenario. Habitat configurations include back, front, or both back and front of the traveling habitat patch, indicating which areas of the habitat are capable of supporting population growth. Environmental scenario describes which transition matrix is used in the back and/or front of the traveling habitat patch: (1) the zero matrix (death outside the patch), (2) no recruitment, (3) no fecundity, or (4) reduced survivorship and growth. In all cases, populations reach a pattern of sustained growth by 20 generations.

## Appendix S4: Critical rates of climate change for supplementary habitat configurations

In the main text, we explore the role of fecundity, recruitment, growth and survival behind the moving habitat patch, with death in front of the habitat patch. However, survival behind the patch and death in front of the patch is just one possible habitat configuration. Survival in front of the patch with death behind the patch is another possible habitat configuration, as well as survival both in front of and behind the moving habitat patch. Here, we explore the role of these other habitat configurations.

Including individuals at the leading edge of the patch increases the estimated critical speed of climate change that a population can endure (fig. S3). These increases are the largest for populations with intermediate dispersal ability. Populations with low dispersal ability do not benefit from having individuals at the leading edge of the patch, regardless of the environmental scenario at the front of the patch. Populations shifting into a habitat where no seed dispersal was possible, but survivorship and growth were at normal levels (scenario 3), had the highest critical rates of climate change. When the habitat in front of the patch caused decreased survivorship and growth (scenario 4), the critical speed of climate change increased relative to the scenario of death in front of the patch (scenario 1). Recruitment failure in front of the patch (scenario 2) produced nearly the same critical rates of climate change as death in front of the patch (scenario 1).

Allowing survival outside of the moving habitat patch (both behind the patch and in front of the patch) increased the critical speed of climate change across all environmental scenarios (fig. S4). When there was no seed production outside the patch, but growth and suvivorship were unchanged (scenario 3), the critical speed of climate change was highest. These high critical speeds of climate change were almost matched when there was no recruitment outside of the patch (scenario 2). Reducing survivorship and growth outside of the patch (scenario 4) provided a moderate increase in the critical speed of climate change compared to the null model of death outside the moving habitat patch (scenario 1). These increases in the critical speed of climate change were greatest for populations with intermediate average dispersal distances, and there was nearly no increase in the critical speed of climate change for populations of poor dispersers.

**Figure S3:**
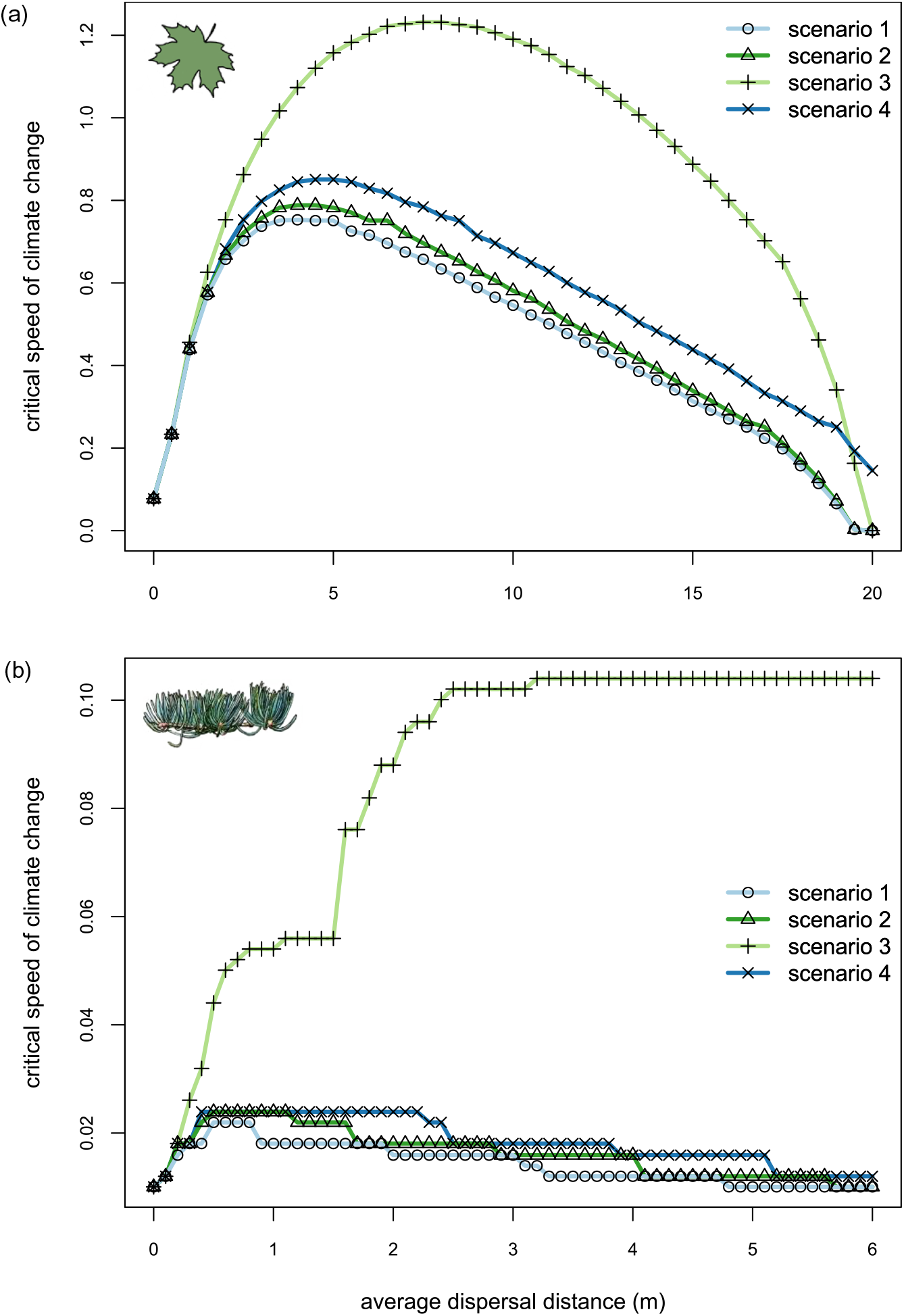
Effect of environment in front of the patch and dispersal distance on the critical speed of climate change for population persistence in sugar maple (a) and white fir (b), with no survival (environmental scenario 1) behind the patch. In front of the patch environmental scenarios tested include no survival (scenario 1), no recruitment (scenario 2), no fecundity (scenario 3), and reduced survivorship and growth (scenario 4).

**Figure S4:**
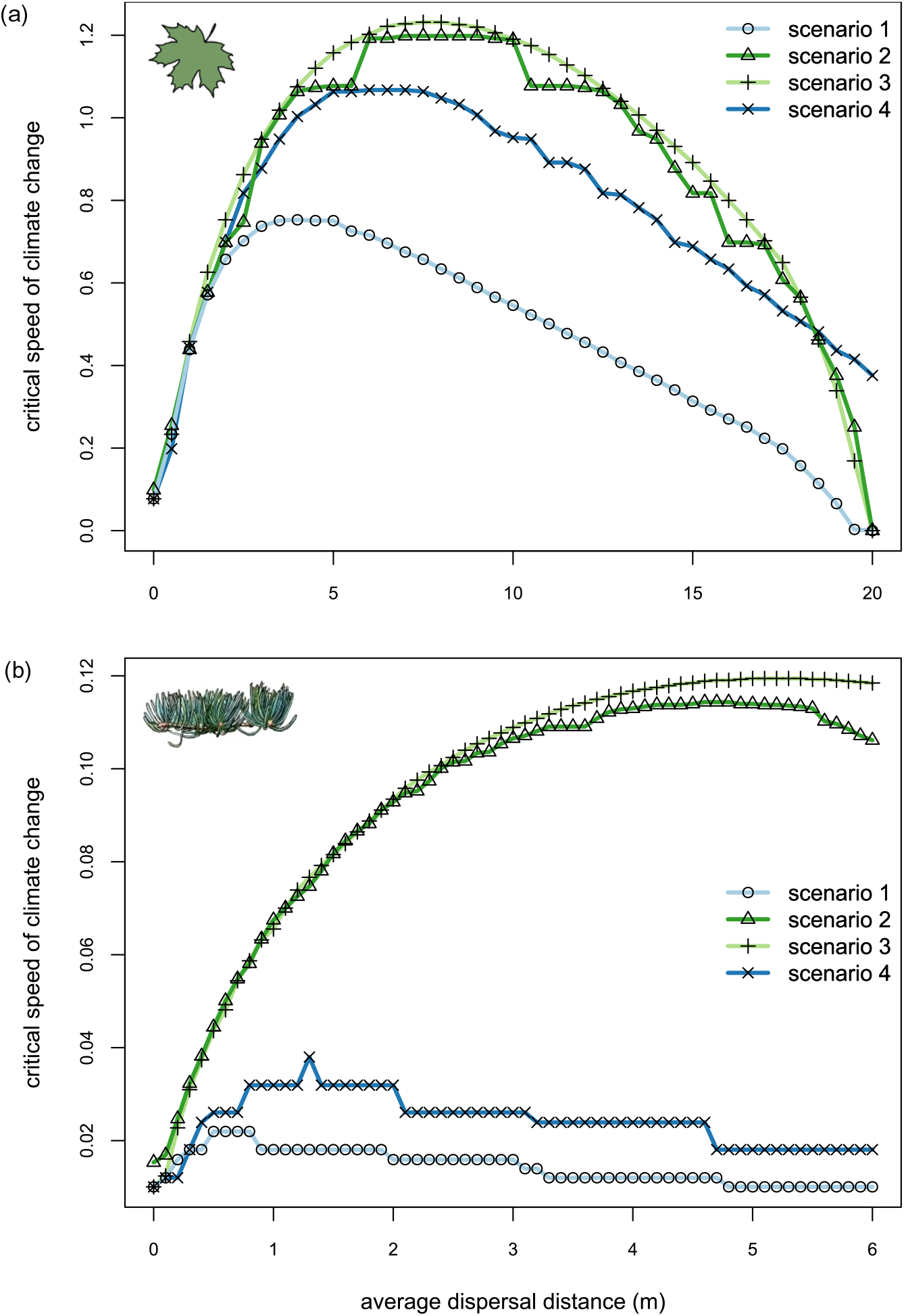
Effect of environment outside of the patch and dispersal distance on the critical speed of climate change for population persistence in sugar maple (a) and white fir (b). Outside of the patch environmental scenarios tested include no survival (scenario 1), no recruitment (scenario 2), no fecundity (scenario 3), and reduced survivorship and growth (scenario 4); in all cases both behind and in front of the patch suitable habitat.

